# PCPE2 and SR-BI Partner to Impact Accumulation of Fat in Mice

**DOI:** 10.1101/298208

**Authors:** Hao Xu, Sushma Kaul, Rachel Kallinger, Michael J. Thomas, Rebecca L. Schill, Daisy Sahoo, Mary G. Sorci-Thomas

## Abstract

LDL receptor knockout mice (LDLr^-/-^) were crossed with PCPE2 knockout mice to obtain *Ldlr*^-/-^,*Pcpe2*^-/-^ mice. The rationale of these studies was to examine the effects of an extracellular matrix protein, PCPE2, on fat storage in a dyslipidemic mouse model. Male *Ldlr*^-/-^, *Pcpe2*^-/-^ mice were fed a Western diet for 25 weeks and their plasma triglyceride metabolism and triglyceride storage was examined. Interestingly, visceral but not subcutaneous fat pad were smaller in diet-fed *Ldlr*^-/-^,*Pcpe2*^-/-^ mice compared to controls. There was no difference in the fatty acid distribution in triglyceride and cholesteryl esters (CE) among the genotypes. *Ldlr*^-/-^, *Pcpe2*^-/-^ mice have higher plasma triglyceride levels and reduced lipoprotein lipase activity. Immunoprecipitation of SR-BI from cell extracts co-precipitated PCPE2 suggesting that PCPE2 and SR-BI are tightly associated. This work also showed that in the absence of PCPE2 SR-BI does not transfer CE from HDL into the cell. These results suggest that HDL, PCPE2, SR-BI, and possibly LPL are associated in an interactome that is required for CE transport into the cell. In the absence of these interactions lipid transport is significantly disrupted.

---

Previous studies have demonstrated a role for procollagen endopeptidase enhancer 2 (PCPE2) in the redistribution of lipids, in particular cholesterol, between tissues cell and plasma(1-6). Therefore, PCPE2 must play a significant role cholesterol homeostasis. Some of the most telling results came from the analysis of high density lipoprotein (HDL) uptake in mice having global knockouts (KO) for both the low density lipoprotein receptor (LDLr) and PCPE2 (*Ldlr*^-/-^, *Pcpe2*^-/-^). HDL isolated from *Ldlr*^-/-^ mice and *Ldlr*^-/-^, *Pcpe2*^-/-^ mice are catabolized at the same rate when injected into *Ldlr*^-/-^ mice. However, *Pcpe2*^-/-^ mice catabolize HDL from *Ldlr*^-/-^ mice and *Ldlr*^- /-^, *Pcpe2*^-/-^ mice at the same rate, but more slowly than *Ldlr*^-/-^ mice(1). These observations, along with others, suggest that the effect of PCPE2 is not at the level of the HDL particle, although particle diameter and composition are affected (1,4), but instead PCPE2 has a more substantial role by directly modulating lipid redistribution and/or uptake.

Curiously PCPE2 is a protein that enhances or facilitates the processing of procollagen to collagen in the extracellular matrix (ECM) (7-9). PCPE2 is related to the enhance protein procollagen endopeptidase enhancer 1, sharing 43% amino acid sequence identity, although, these proteins have different tissue distributions with PCPE2 more abundant in heart, aorta, and adipose (9). Because of its known relationship with collagen processing we reported that the collagen content of the aorta and liver were reduced in the *Ldlr*^-/-^, *Pcpe2*^-/-^ mouse. Preliminary studies indicated that the absence of PCPE2 primarily affected the function of scavenger receptor class BI (SR-BI) (1), the most physiologically relevant HDL receptor. The following studies enlarge our understanding of how these extracellular components participate in cholesterol homeostasis.

PCPE2 is a glycoprotein that has several well-defined protein motifs. The first are a pair of CUB (Complement C1r/C1s, Uegf, Bmp1) domains at the N-terminus separated by a short linker region with each CUB characterized by β- sandwich folds and Ca^+2^ binding sites (10,11). The C-terminal region carries a netrin-like domain that can bind to heparan sulfate proteoglycans (12) located in the ECM, such as glypicans that are associated with the cell membrane by a glycosyl-phosphatidylinositol

## Mouse Studies

### RNA Seq analysis of Ldlr^-/-^ and Ldlr^-/-^ Pcpe2^-/-^ mice

Messenger RNA was isolated from visceral adipose from 3 mice of each genotype using Trizol according to manufacturers’ instructions. RNA Seq was performed on visceral adipose tissue mRNA isolated from Western diet-fed gender- and age-matched PCPE2 knockout (n=4) and control mice (n=4). Libraries were prepared, and sequencing performed on the HiSeq 2500 using TruSeq v4 SBS chemistry and 2×125-paired end-reads. Upon completion of sequencing, reads were demultiplexed to bioinformatically separate samples by index, and aligned using the Illumina Bcl2fastq program. The results were compiled and searched for genes with either up regulated or down regulated, Fig. 1A. As previously reported, *Ldlr^-^* anchor possibly integrated into lipid rafts (13-15). Studies into the mechanism of action of the homolog PCPE1 have suggested that it binds to the C-propeptide region of procollagen III to enhance bone morphogenetic protein 1 activity (16). Using surface plasmon resonance Zhu et al. has suggested that PCPE2 binds to apolipoprotein A-I (apoA-I) of HDL (3), and it is this interaction that might suggest a mechanism for PCPE2 modulation of SR-BI activity.

**Figure 1.**
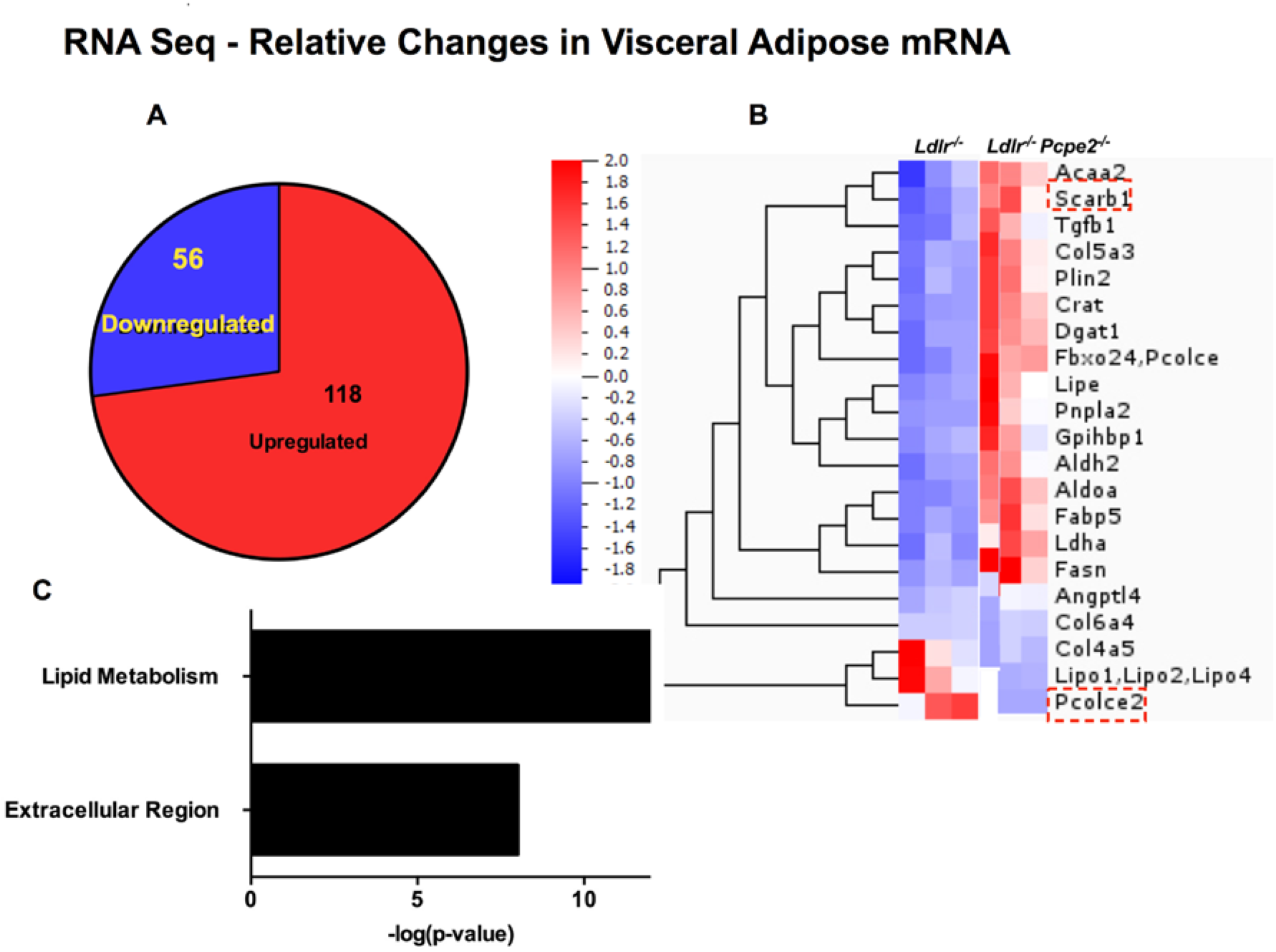
RNA Seq analysis showing relative changes in visceral adipose mRNA. mRNA was isolated from visceral adipose tissue of *Ldlr^-/-^* and *Ldlr^-/-^ Pcpe2^-/-^* mice and then analyzed using RNA Seq. **A** shows the number of up or down regulated genes. **B** is a heat map showing a subset of genes that were regulated and the relative degree of regulation. **C** shows that genes for proteins involved in lipid metabolism or the extracellular region of the cells are highly represented. mRNA for the principal genes under study are outlined in red dotted boxes; *Pcole2*, protein PCPE2, and *Scarb1*, protein SR-BI. There are 3 mice in each group.

This communication provides new experimental evidence for the role of PCPE2 in influencing SR-BI activity in adipose tissue. Several lines of analysis have suggested that PCPE2 and SR-BI are associated on the cell surface, although, they may not be in direct contact. More results are provided showing that in the absence of PCPE2 delivery of cholesteryl esters (CE) from HDL into adipocytes is substantially reduced. The effect of a global knockout of PCPE2 on mice (*Ldlr*^-/-^, *Pcpe2*^-/-^) was to increase the size of fat droplets in visceral adipose tissue (VAT) and in subcutaneous fat (SAT) compared to the same fat depots in *Ldlr*^-/-^ mice, however, with a total reduction in fat mass. There were no apparent differences in connective tissue and tissue macrophage populations between knockout mice and the *Ldlr*^-/-^ background.

## Results

^*/-*^ *Pcpe2^-/-^* mice expressed higher levels of SR-BI protein(1) consistent with the higher levels of *Scarb1* mRNA found by RNA Seq analysis, Fig. 1B. In contrast, *Pcolce2* and PCPE2 mRNA levels were elevated in *Ldlr^-/-^* mice, while the levels of *Scarb1* mRNA were reduced. The analysis also showed that genes associated with lipid metabolism and extracellular proteins were more highly expressed in *Ldlr^-/-^ Pcpe2^-/-^* mouse adipose, Fig. 1C.

### Fat depots in Ldlr^-/-^ and Ldlr^-/-^ Pcpe2^-/-^ mice

*Ldlr^-/-^* and *Ldlr^-/-^ Pcpe2^-/-^* mice were fed chow or a Western diet for 25 weeks, weighed, and then sacrificed by Institutional-approved protocols. Fat depots were removed and weighed. Although body weight was not statistically different between the two genotypes, the amount of visceral fat was less in *Ldlr^-/-^ Pcpe2^-/-^* mice, while subcutaneous and brown fat (data not shown) were not statistically different between genotypes, Fig. 2A-C. Microscopy of H & E stained sections, Fig. 2D, showed that the visceral and subcutaneous fat from *Ldlr*^-/-^*Pcpe2*^-/-^ mice had fewer, but larger cells than did *Ldlr^-/-^* mice. This was backed up by cell surface counting of cell size range that showed *Ldlr^-/-^* adipose had significantly more cells in the 500 to 2000 µm^2^ range than did *Ldlr*^-/-^*Pcpe2*^-/-^ adipose cells, which displayed a much broader range of sizes (data not shown).

**Figure 2.**
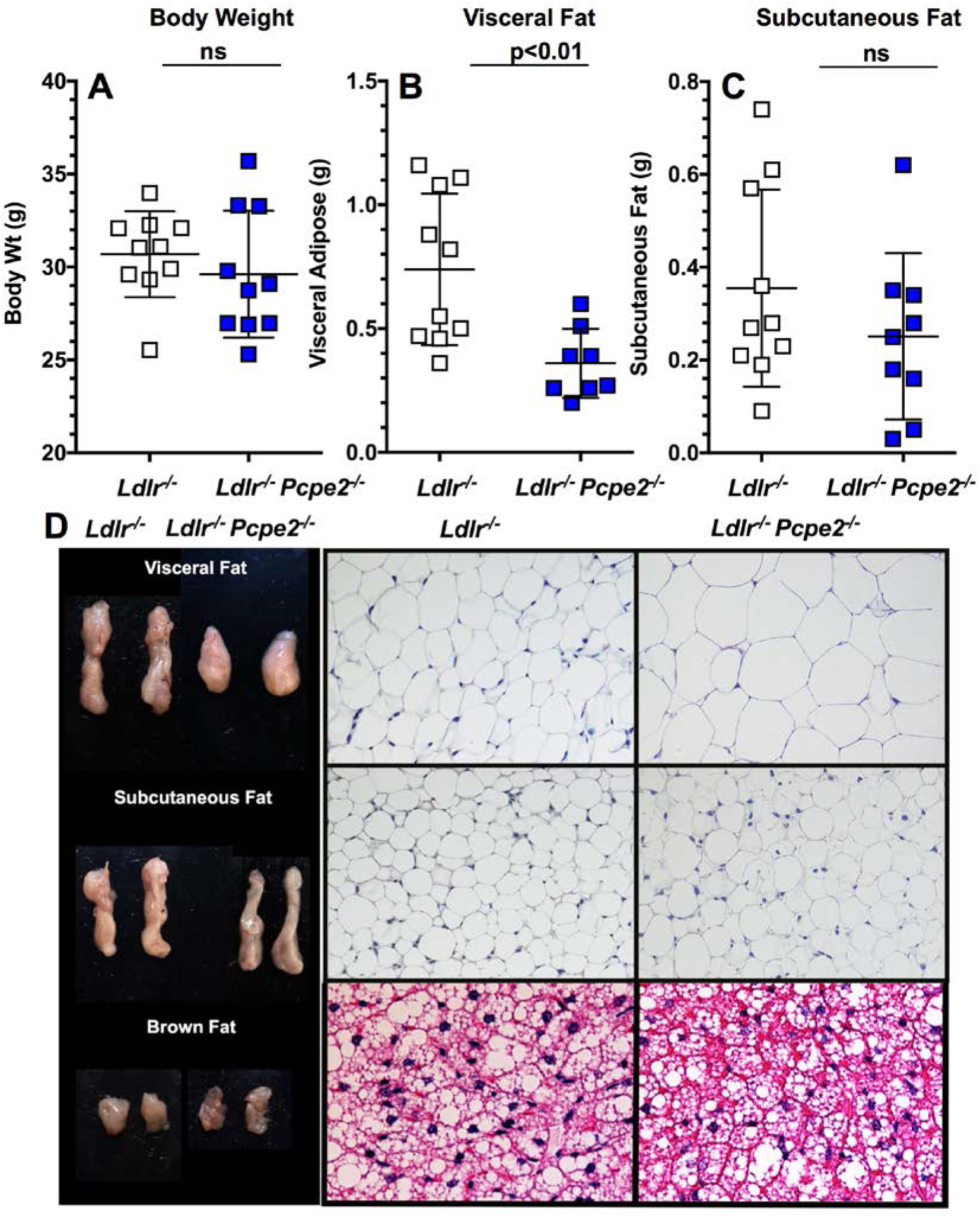
Body weight and adipose tissue from *Ldlr^-/-^* and *Ldlr^-/-^ Pcpe2^-/-^* mice. **A**. The upper left panel shows the body weight of mice (10/Group) fed a Western diet for 25 weeks. **B** and **C**. The next two panels show the mass of visceral fat and subcutaneous fat, n = 8-10, removed from these mice after 12 weeks on a western diet. **C**. The left panel shows typical depots of each type of fat while the right panel shows typical H & E stains for each of the fat depots from each genotype.

### Lipase activity in Ldlr^-/-^ and Ldlr^-/-^ Pcpe2^-/-^ mice

Previous studies had demonstrated that *Ldlr^-/-^ Pcpe2^-/-^* mice had greater concentrations of cholesterol and triglyceride (TG) than did *Ldlr^-/-^* mice(1). These differences were accentuated on the Western diet. Additional studies were performed to establish how hepatic lipase and lipoprotein lipase influence the concentration of plasma TG. Fig. 3A shows that *Ldlr^-/-^ Pcpe2^-/-^* mice produce triglyceride at a faster rate than do *Ldlr^-/-^* mice after blocking TG hydrolysis with Triton. In Fig. 3B lipoprotein lipase and hepatic lipase activity were measured in *Ldlr^-/-^* and *Ldlr^-/-^ Pcpe2^-/-^*. In all cases the lipase activity of *Ldlr^-/-^ Pcpe2^-/-^* mice was lower than the *Ldlr^-/-^* controls.

**Figure 3.**
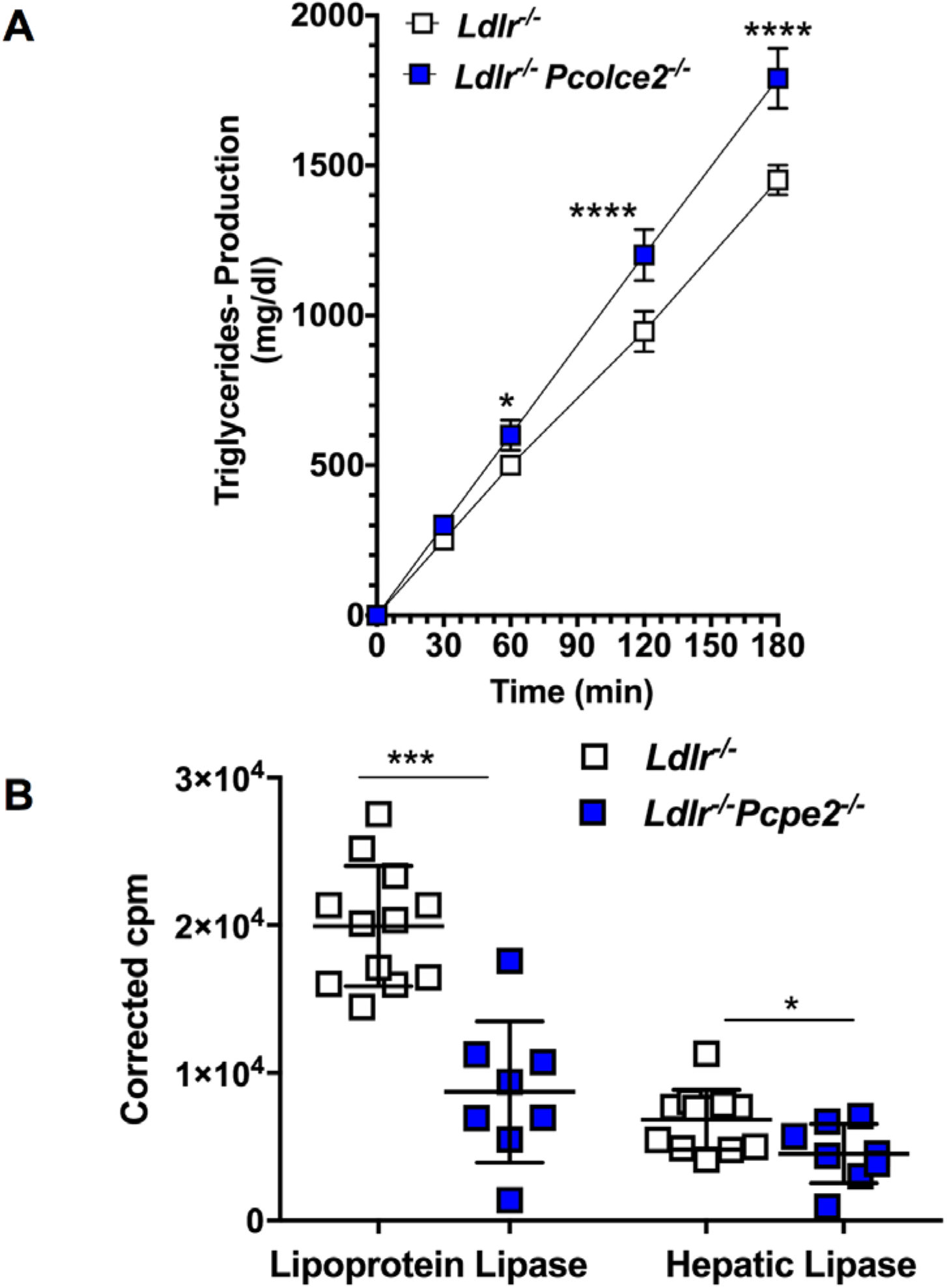
Activities of triglyceride lipases. **A** shows the total production of triglyceride in the plasma of *Ldlr^-/-^* and *Ldlr^-/-^ Pcpe2^-/-^* mice after triton block (5 mice/group). **B.** Lipase activity was measured in plasma from *Ldlr^-/-^* (12) and *Ldlr^-/-^ Pcpe2^-/-^* (8) mice at 30 min. Plasma triglyceride concentrations were determined and triglyceride production was calculated after subtracting basal values. Mean ± SD.* p<0.05, **p<0.01, ***p<0.001.

### Cholesterol levels in liver and adipose tissue

Liver and adipose tissue was extracted and analyzed by GC-MS/MS for free cholesterol (FC) and total cholesterol (TC). Fig. 4A shows that *Ldlr*^-/-^*Pcpe2*^-/-^ mice had less FC and EC per mg wet liver than did *Ldlr^-/-^* mice. In Fig. 4B it is evident that in VAT tissue FC levels were reversed with *Ldlr*^-/-^*Pcpe2*^-/-^ mice having more FC, but with less CE in *Ldlr*^-/-^*Pcpe2*^-/-^ mice. The *Ldlr*^-/-^ mouse liver has about 20-fold more FC/mg tissue than adipose tissue and about 30- fold more EC/mg tissue. For *Ldlr*^-/-^*Pcpe2*^-/-^ mice the ratios were 10-fold and 25-fold, respectively. The distribution of fatty acids (FA) in liver was the same for both genotypes, 18:1>16:0>16:1>18:0>20:1>14:0>18:2. VAT tissue the ratios were different compared to liver tissue and somewhat different between the genotypes. For *Ldlr*^-/-^*Pcpe2*^-/-^ mice the ratio was 18:1>>16:0>16:1~18:2>18:0>20:4, while for the *Ldlr*^-/-^ mice the ratios were 18:1>>16:0>16:1>18:2>18:0~20:4. Liver CE carried about 71% monounsaturated FA and 1% polyunsaturated FA, while VAT tissue had about 62% monounsaturated FA and 13% polyunsaturated FA. The 16:1/16:0 ratio for liver was 0.78 for *Ldlr*^-/-^ mice and 0.63 for *Ldlr*^-/-^ *Pcpe2*^-/-^ mice. The same ratio for VAT was 0.9 and 0.83, respectively.

**Figure 4.**
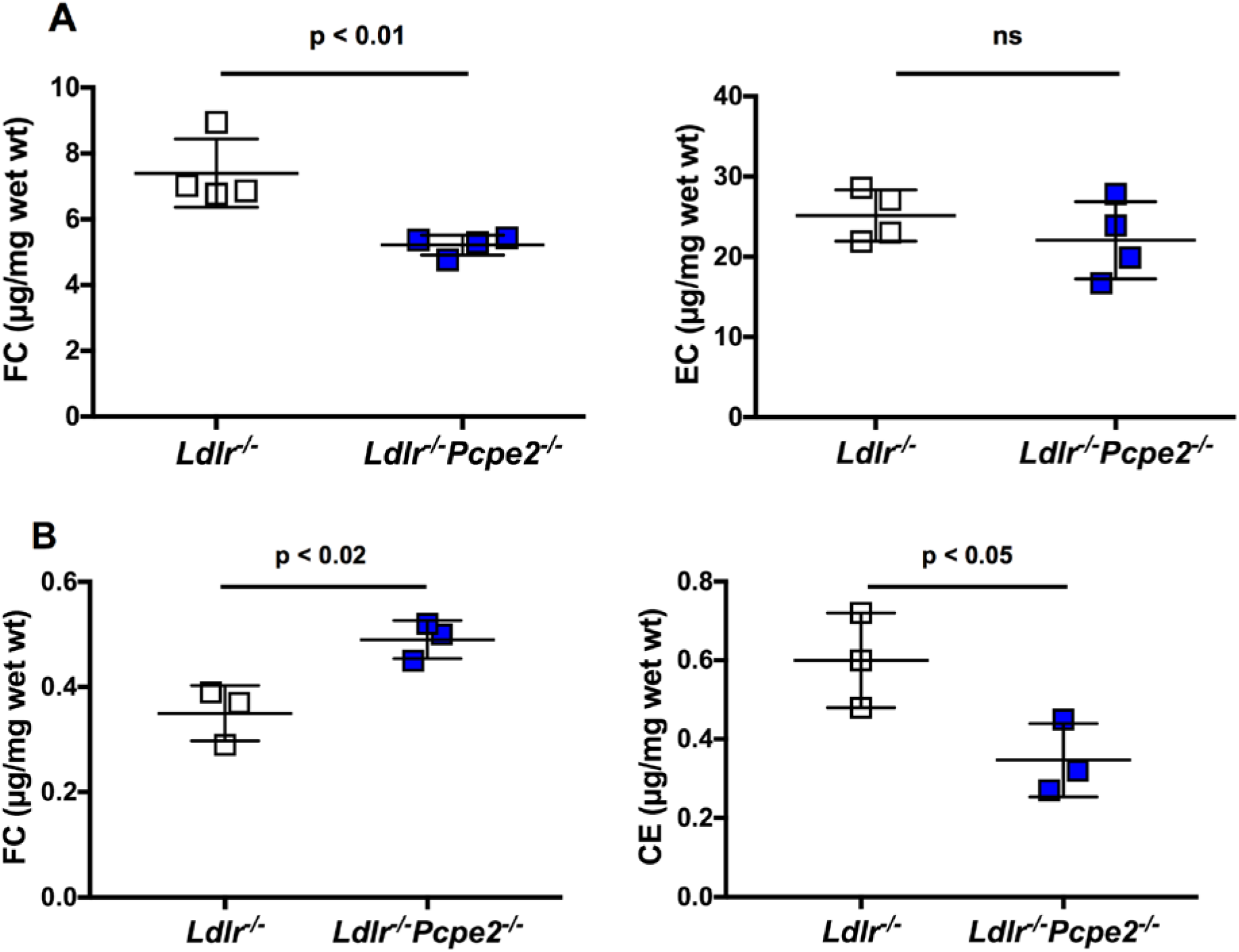
Cholesterol and cholesteryl ester in liver and VAT tissue. Samples of liver and VAT tissue from *Ldlr*^-/-^ and *Ldlr*^-/-^ *Pcpe2*^-/-^ mice were extracted and then analyzed by GC-mass spectrometry for free cholesterol in liver, **A**, and adipose tissue, **B**. Mice had been maintained on a Western diet for 25 weeks. Samples were independently analyzed by LC-mass spectrometry for liver and adipose cholesteryl esters. There were three or more mice per group.

### Triglyceride levels in liver and adipose tissue

Liver and adipose tissue was extracted and analyzed by LC-MS/MS for TG. Fig. 5 shows that *Ldlr*^-/-^ and *Ldlr*^-/-^*Pcpe2*^-/-^ mice had about the same amount of TG per mg wet liver. VAT tissue also showed no statistical difference on the amount of TG per mg wet tissue. VAT tissue had about 30-fold more TG than did liver. Liver had the following distribution of fatty acids for both genotypes; 18:1>16:0>16:1>18:0>20:1>14:0>18:2, while VAT tissue had the following distribution of fatty acids for both genotypes; 18:1>16:0>16:1>14:0≈18:2>18:0.

**Figure 5.**
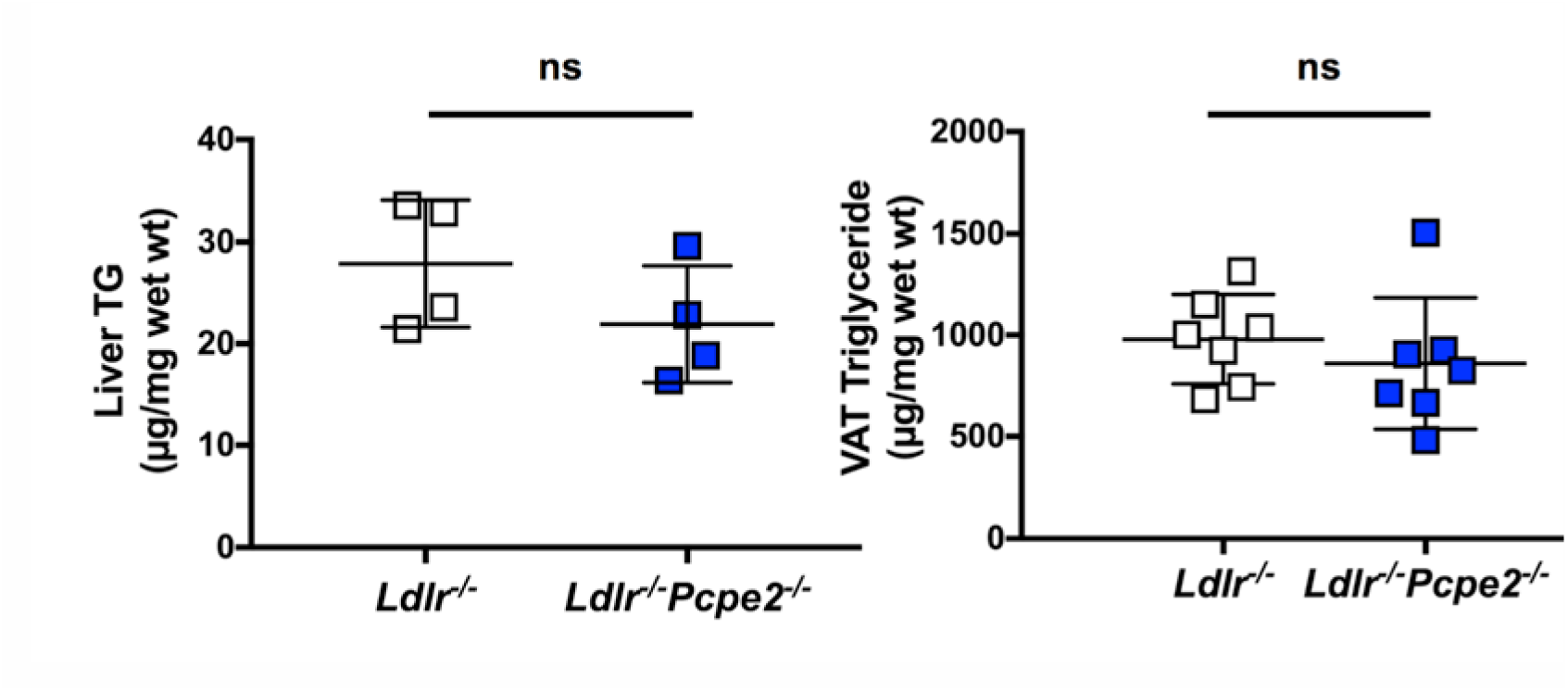
Triglyceride in liver and VAT adipose tissue. Samples of liver and VAT tissue from *Ldlr*^-/-^ and *Ldlr*^-/-^ *Pcpe2*^-/-^ mice were extracted and then analyzed by LC-mass spectrometry for triglyceride. Mice had been maintained on a Western diet for 25 weeks. There were four or more mice per group. The means of the groups were not significantly different from one another.

## Cell Studies

Fig. 6 extends our published findings by presenting results that circumvent issues related to overexpression of PCPE2 or SR-BI and their partnership. Using CRISPR/Cas9 methodology Chinese hampster overy (CHO) cells were generated that lacked PCPE2. CHO cells were chosen for these proof-of-concept studies because they express high levels of both SR-BI and PCPE2, Fig. 6A, providing a direct test of the effect of PCPE2 on SR-BI function. To validate our premise, we also created 3T3 cells, which normally express both PCPE2 and SR-BI, Fig. 6C, that lack PCPE2 and compared their ability to remove HDL-cholesteryl ester (CE) with wild-type 3T3 cells. Figures 6B and 6D show that CHO^*Pcpe2*-/-^ cells had a 20% reduction in CE uptake compared to CHO^*WT*^ cells, while 3T3^*Pcpe2*-/-^ cells had a 47% reduction compared to 3T3^*WT*^ cells. These reductions in CE transport are taking place with about a 10-fold increase of SR-BI in each cell line when PCPE2 had been knocked out.

**Figure 6.**
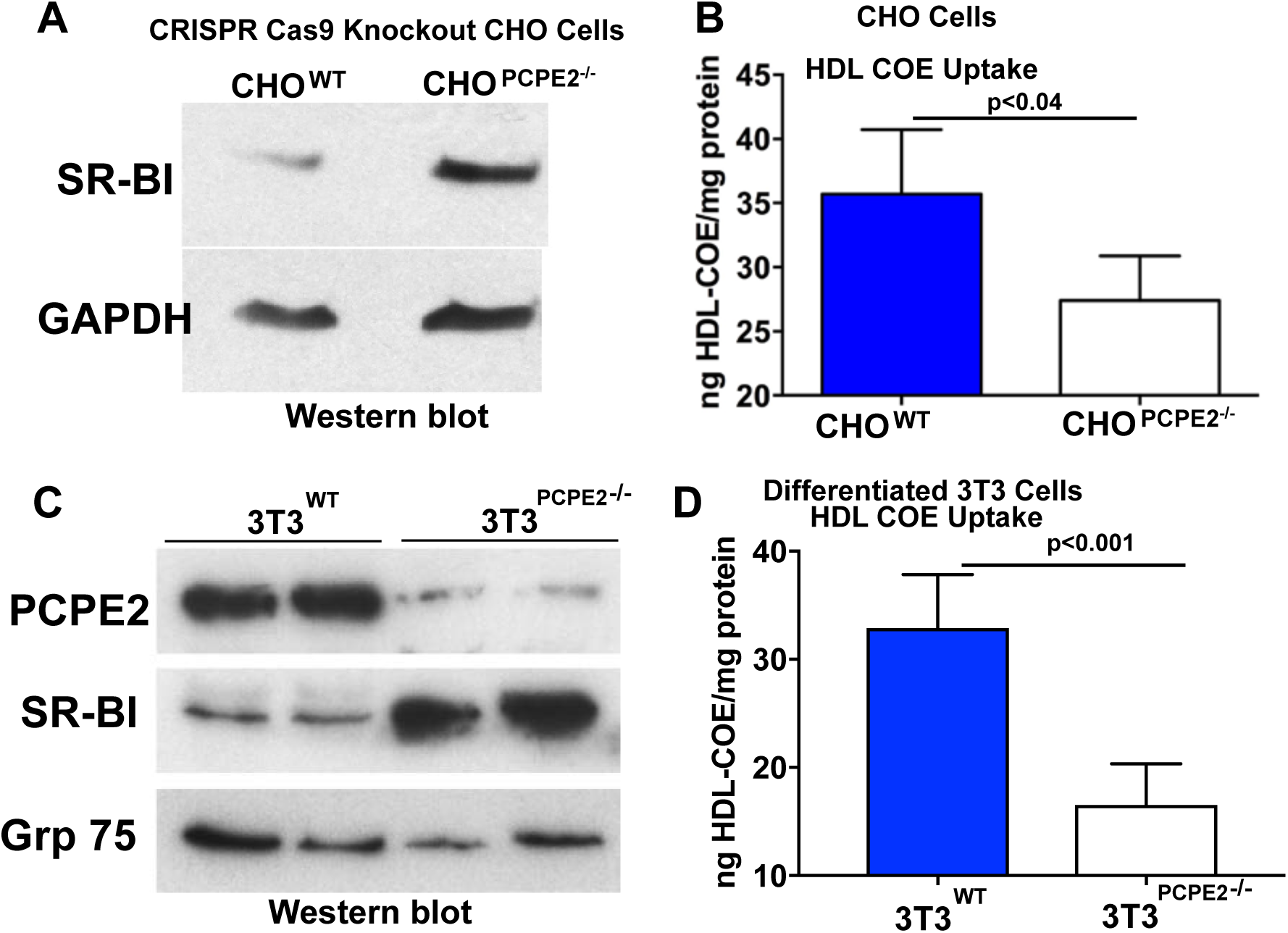
Reduced SR-BI Function Despite Increased Protein Expression Associated with the Loss of PCPE2. **A)** Increased SR-BI protein expression in CHO^PCPE2-/-^ cells. **B)** Uptake of ^3^H-cholesteryl- oleoyl-ether (COE)-labeled HDL was reduced in CHO^PCPE2-/-^ cells compared to CHO^WT^ cells. **C)** Western blot of differentiated 3T3 cell total protein extracts probed for PCPE2, SR-BI and Grp75 (load control) after 9 days. **D)** Uptake of ^3^H-COE-labeled HDL was reduced in differentiated 3T3^PCPE2-/-^ compared to 3T3^WT^ cells.

### Lipid accumulation in 3T3 cells

Human and mouse adipocytes contain relatively large amounts of PCPE2 and SR-BI. 3T3 cells can be differentiated to adipocyte-type cells over several days (17). These differentiated cells begin to accumulate lipid by day 5, Fig. 7A, as indicated by the increased staining with oil red O. Panel B shows that both SR-BI and PCPE2 were up regulated by day five with protein levels that increased or remained constant to day 9. More careful examination of 3T3^*WT*^ and 3T3^*Pcpe2*-/-^ cells after differentiation followed by treatment with Oil Red O shows that wild-type cells have considerably more lipid droplets and perilipin than 3T3^*Pcpe2*-/-^ cells (data not shown).

**Figure 7.**
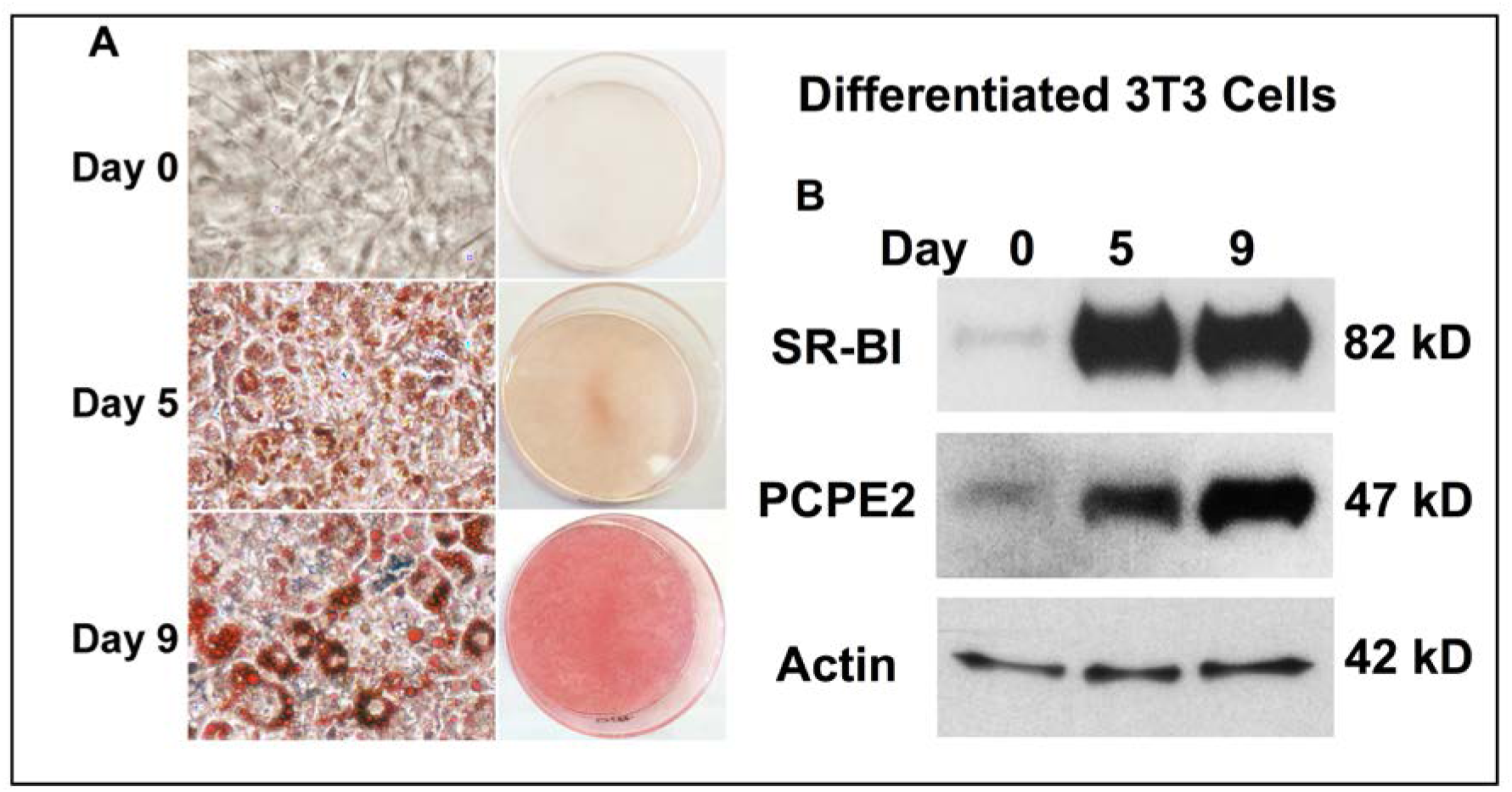
Differentiation of 3T3 cells to adipocyte-type cells. 3T3 cells were plated and grown to confluence, then allowed to differentiate for 9 days. **A)** photographs at days 0 through 9 for cells treated with Oil-Red O to accentuate lipid accumulation. Closeup photomicrographs were recored at X10. **B)** A Western blot for SR-BI, PCPE2, and actin (load control) from equal amounts of protein separated by SDS-PAGE. Note that both SR-BI and PCPE2 were upregulated by day 5.

### Consequences of the interaction of PCPE2 and SR-BI

Our results indicate that there is a strong association between SR-BI and PCPE2. In the first set of studies whole cell protein extracts were analyzed by nondenaturing gradient gel electrophoresis (NDGGE) with and without the mild detergent perfluorooctanoic acid (PFO). 10 µg of protein extract were analyzed and the gel probed with SR-BI antibody (Norvus, NB400-Earlier studies had suggested that the absence of PCPE2 had significant consequences for the transport of cholesteryl ester from HDL into a cell by SR-BI. This report extends these studies, reporting the effect on mice that carry a global knockout of PCPE2. One of the principal changes was found to be in adipose tissue where VAT was substantially reduced in *Ldlr*^-/-^*Pcpe2*^-/-^ mice. Other fat depots showed lesser changes. Overall these animals had higher HDL levels, consistent with our previous reports that SR-BI activity is markedly reduced in the PCPE2 knockout mouse.

Figure 1 shows RNAseq analysis of adipose tissue from *Ldlr*^-/-^*Pcpe2*^-/-^ mice compared to *Ldlr*^-/-^ mice with three mice analyzed in each group. mRNA quality was assessed for each mouse and found to have no detectible degradation. The results showed that 118 genes were up regulated while 56 were down regulated. Criteria for regulation were that the – log(p-value) was greater than +1 or −1. The greatest numbers of genes affected were those associated with lipid metabolism and the extracellular region of the cell. Previous studies had shown that *Ldlr*^-/-^*Pcpe2*^-/-^ mice had high higher plasma cholesterol levels than *Ldlr*^-/-^ mice 101). As shown in the upper panel of Fig. 8 the amount of SR-BI was greater in the extract from 3T3^*Pcpe2*-/-^ cells, but SR-BI dimer was reduced in 3T3^*Pcpe2*-/-^ cells compared to wild-type cells, lower left panel, and completely disappeared, showing up as monomer, when the same concentration of protein was analyzed in a buffer containing 1.7% PFO, lower right panel. In a second set of experiments differentiated 3T3 cells were extracted and the protein extracts treated with an antibody to SR-BI. Fig. 9 shows that PCPE2 was co-precipitated with SR-BI. Fig. 8 showed that the amount of SR-BI increased in 3T3^*Pcpe2*-/-^ cells compared to wild-type 3T3 cells. Identical results were obtained when wild-type CHO cells were treated with small interfering RNA to PCPE2 and in protein extracts from CHO^*Pcpe2*-/-^ cells prepared using CRISPR/Cas9 methods (Data not shown). These observations taken together suggest that cells may be able to sense the absence of SR-BI activity and attempt to synthesize additional SR-BI to compensate.

**Figure 8.**
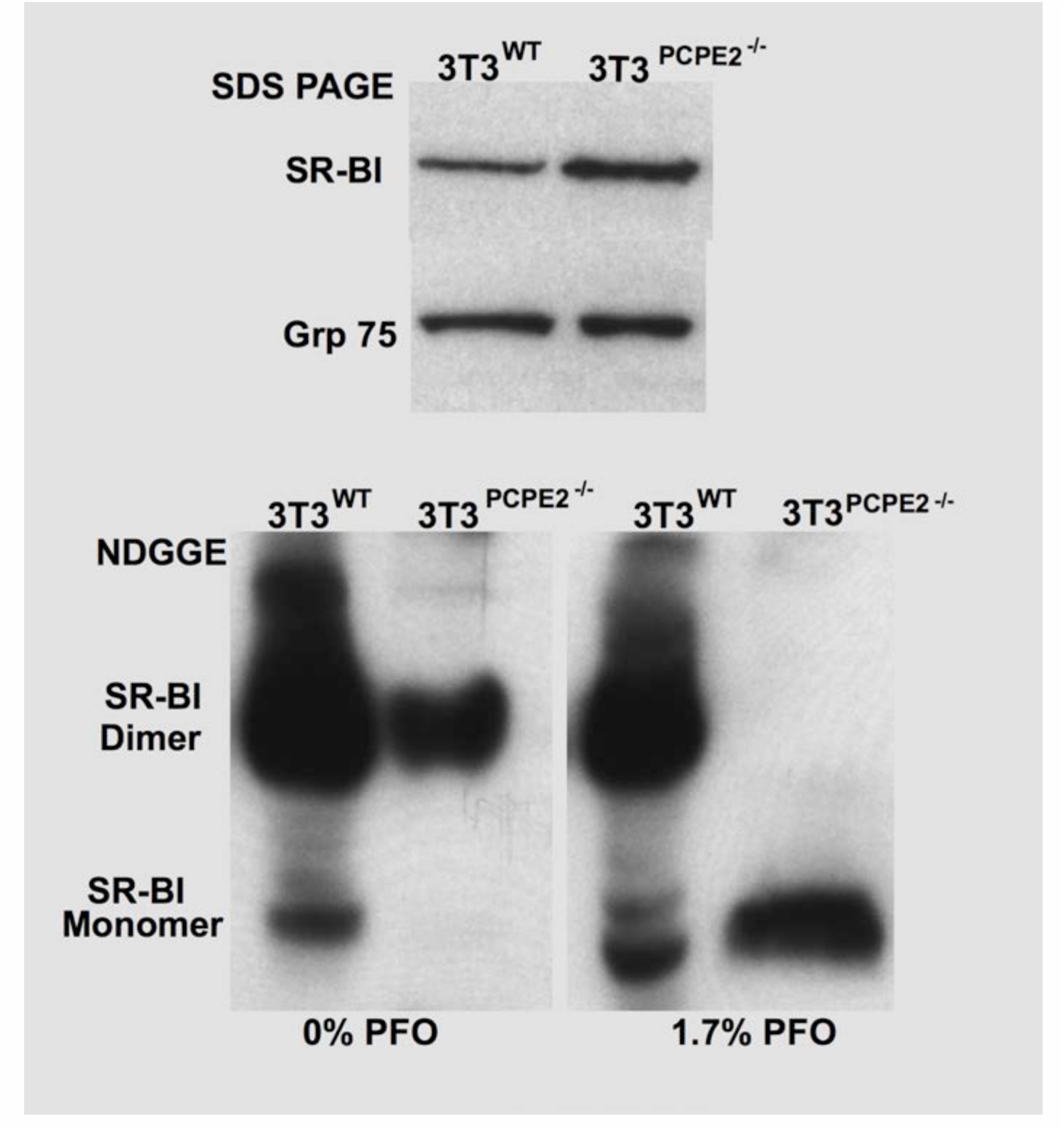
PCPE2 strengthens the self-association of SR-BI. The upper panel shows SR-BI levels in equal concentrations of protein extract from 3T3^wt^ and 3T3^*Pcpe2*-/-^ cells analyzed by SDS PAGE and developed with SR-BI antibody (Norvus, NB400-101). The PCPE2 KO cells had noticeably more SR-BI than the wild-type cells. In the lower left panel is shown the wild-type and 3T3^*Pcpe2*-/-^ extracts analyzed by NDGGE, which should be compared to the lower right panel that includes the mild detergent PFO. These lower panels show that the SR-BI is more tightly associated in wild-type cells than in 3T3^*Pcpe2*-/-^ cells.

**Figure 9.**
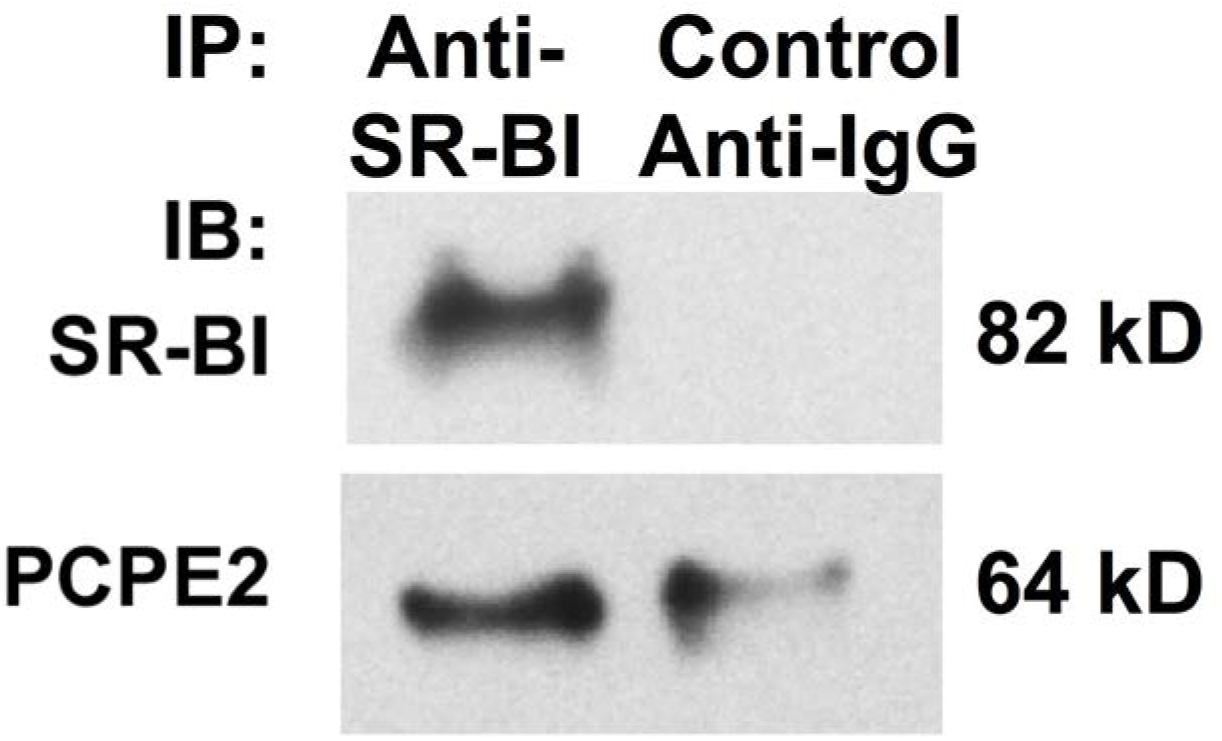
Antibodies to SR-BI also precipitate PCPE2. 3T3 cells are differentiated, a protein extract was prepared, and the extract treated with antibody to SR-BI (Norvus, NB400-101). The precipiate was analyzed using SDS PAGE and the individual regions analyzed by Western blot analysis using antibodies to SR-BI(Norvus, NB400-101) and PCPE2 (Abcam, ab156224).

## Discussion

indicating that cholesterol was less effectively metabolized when PCPE2 was absent. As anticipated *Pcole2* was down regulated in *Ldlr*^-/-^ *Pcpe2*^-/-^ mice with respect to the *Ldlr*^-/-^ and as previously reported from protein immunoblots of mouse liver (1) *Scarb1* was up regulated in *Ldlr*^-/-^*Pcpe2*^-/-^ mice compared to *Ldlr*^-/-^ mice. Although not explored in this manuscript, the increased levels of SR-BI suggest, but don’t prove, that cholesterol transport by SE-BI is essential for optimal cellular function.

When *Ldlr*^-/-^*Pcpe2*^-/-^ and *Ldlr*^-/-^ mice are fed a Western diet the total body weights for both genotypes are comparable within experimental error. However, analysis of the body fat shows that *Ldlr*^-/-^*Pcpe2*^-/-^ visceral fat pads were about 50% smaller than those from *Ldlr*^-/-^ mice. However, the amount of visceral fat TG per mg wet weight was the same for both genotypes. Other fat depots did not show a significant change in the fat pad mass between the two genotypes. However, the adipose cells in all white fat depots from *Ldlr*^-/-^*Pcpe2*^-/-^ mice were larger than cells from *Ldlr*^-/-^ mice. Previous studies suggested showed that plasma TG levels were twice as high in Western diet-fed *Ldlr*^-/-^ *Pcpe2*^-/-^ mice compared to *Ldlr*^-/-^ mice (1). TG accumulation in plasma was about 30% faster in *Ldlr*^-/-^*Pcpe2*^-/-^ mice consistent with decreased activity of both lipoprotein lipase and hepatic lipase in *Ldlr*^-/-^*Pcpe2*^-/-^ mice or suggesting that uptake by other lipoprotein receptors was reduced. Increased lipoprotein lipase activity has been associated with obesity and insulin resistance (18) and in adipocytes is usually associated with reduced glucose metabolism (19). Interestingly, specifically knocking out lipoprotein lipase in adipose tissue increased plasma TG, but had minimal effects on white fat mass, affecting brown fat metabolism to a much greater extent (20).

The knockout of *Pcole2* reduces PCPE2 levels and is associated with reduced SR-BI transport of CE in livers of *Ldlr*^-/-^*Pcpe2*^-/-^ mice (1). To study SR-BI related CE transport a CHO was generated with *Pcole*2 silenced. This cell line showed reduced uptake of the CE analogue cholesteryl-oleoyl-ether (COE), as was reported for *Ldlr*^-/-^*Pcpe2*^-/-^ mice. Protein extracts from mouse liver and RNAseq analysis of adipose tissue also showed that SR-BI levels are dramatically increased in the absence of PCPE2. To study the mechanism of transport involving the SR-BI/PCPE2 interaction in adapose tissue a 3T3 cell line was developed using CRISPR/Cas 9 knockout of PCPE2. The 3T3^PCPE2-/-^ cell line, like the 3T3^WT^, was shown to differentiate into adipocyte-like cells that accumulated lipid droplets with an increased production of SR-BI and PCPE2, Fig. 6. Like has been observed for other cells and tissues, the 3T3^PCPE2-/-^ cell line showed substantially increased SR-BI levels compared to 3T3^WT^, but a decreased ability to internalize COE.

The dependence of SR-BI activity on PCPE2 suggests that there might be interaction between the proteins. This interaction could take at least two forms. The first is that PCPE2 promotes or stabilizes the oligomerization of SR-BI, a step that has been proposed as critical SR-BI promoted CE transport (21-23) and then SR-BI oligomers bind HDL. Figure 8 provides evidence that SR-BI may exist as a dimer in wild type cells. Protein extracts from 3T3^wt^ cells show a strong dimer band when analyzed using Standard RNA extraction and purification techniques performed followed by cDNA library synthesis (TruSeq Stranded mRNA LT Set B, nondenaturing gradient gel electrophoresis, while 3T3^*Pcpe2*-/-^ extracts show a much weaker dimer band. When the samples were run in the presence of the mild detergent perfluorooctanoic acid the wild type sample retained the dimer as the predominant form of SR-BI while the extract from 3T3^*Pcpe2*-/-^ cells shows only monomer. When these samples were immunoprecipitated with anti-SR-BI and the precipitate analyzed by SDS PAGE both SR-BI and PCPE2 were found by Western blotting. The second interaction mode may require HDL with PCPE2 binding HDL, which then binds to SR-BI. Apolipoprotein AI, the principal protein component of HDL, has been reported to bind PCPE2 (3).

Summarizing the results of these studies the absence of PCPE2 SR-BI dimerization is reduced and the transport of CE is substantially reduced. Physiologic effects include, reduced VAT, reduced activity of lipoprotein lipase, increased plasma TG and TC, cholesterol excretion was substantially reduced after the loss of PCPE2 (1) and increased HDL size. Previous studies demonstrated that the increase in particle size and composition was not responsible for changes in lipid metabolism (1). These results suggest that PCPE2 is an essential component of a local set of protein interactions, called an interactome, that is located around SR-BI in specialized lipid rafts called caveolae (24,25). This interactome of PCPE2/SR-BI/HDL may also include lipoprotein lipase and the adaptor protein, PDZ domain containing 1 (PDZK1), which associates with SR-BI through its C-terminal cytoplasmic leg (26,27) and couple it to Src kinase(s) (28). In endothelial cells this interactome is associated with nitric oxide release (29,30) that involves the participation of several additional kinases and requires the transport of CE by SR-BI. What may be unusual about this local interactome is that it spans from the plasma, HDL, into the extracellular matrix, PCPE2, through the membrane, SR-BI, to a cytoplasmic protein, PDZK1.

## Experimental procedures

### RNA Seq Analysis

Illumina). The quality of the cDNA libraries was assessed using an Agilent 2100 Bioanalyzer and Agilent DNA 1000 kit. Final quantification of prepared libraries was performed using qRT-PCR and MiSeq 50. RNA sequencing was performed on the HiSeq 2500 using TrusSeq v4 SBS chemistry and 2×125 paired-end reads. Primary RNAseq analysis was performed through an analytical pipeline consisting of Bowtie and Tophat for read mapping and alignment, Cufflinks for transcript assembly, Cuffquant for transcript abundance quantification, Cuffnorm for normalization, and Cuffdiff for differential expression. Gene- or isoform-level transcript abundances were quantified as fragments per kilobase per million reads. Ingenuity Pathway Analysis was used to analyze genes exhibiting significant expression alterations, with a false discovery rate of <0.026 compared to wild type.

### In Vivo TG Secretion

In vivo hepatic TG secretion was measured after detergent block of TG lipolysis (31). Briefly, male and female mice (5/group) were anesthetized and injected in the peritoneal cavity with Triton WR-1339, 500 mg/kg, to block lipolysis and [^3^H]oleate (5 µCi/g body weight) to trace TG synthesis.

### Triglyceride Lipase Assay

These analyses were kindly performed by Dr. Goldberg’s group using the method described in Garcia-Arcos et al.(20). Briefly (32), to measure hepatic TG production rate, post-heparin plasma was drawn from fasted mice 5 min after tail vein injection of 100 units of heparin/kg body weight. Plasma was mixed with 10% Intralipid/[^3^H]TG emulsion as substrate with human serum for apolipoprotein CII. Hepatic lipase activity was determined by including 1 mM NaCl in the solution. Lipoprotein lipase activity was determined by subtracting the hepatic lipase numbers from the total lipase activity.

### Animals and Diet

*Ldlr*^-/-^ and *Ldlr*^-/-^ *Pcpe2*^-/-^ mice (1) were placed on a high fat diet (42% calories from fat, 0.2% calories from cholesterol) (Envigo-Teklad, #TD 88137) at 4 weeks of age. Mice were fasted for 3 hours prior to being anesthetized with ketamine/xylazine. Euthanasia and blood collection were by cardiac puncture. Numbers of specific genotype used for each study/assay are indicated in each of the figure legends. Mice were bred and housed in microisolator cages in a pathogen-free animal facility at the Medical College of Wisconsin (MCW). All experiments followed guidelines of the MCW Animal Care and Use Committee and the approval of the use of rodents was obtained from MCW according to criteria outlined in the Guide for the Care and Use of Laboratory Animals from the National Institutes of Health.

### H & E Staining

The Pathology Histology Core Facility of MCW performed sectioned adipose tissue and performed H & E staining. Sections were digitized using a Nikon microscope and Image-Pro Plus 6.2 software. Results were quantified using NIS-Elements software.

### Creation of 3T3^PCPE2-/-^ Cells Using CRISPR/Cas9

Guide RNAs targeting the murine *Pcolce2* gene were cloned into a px459 vector. Plasmid transfected 3T3-L1 cells were enriched by 2 days of puromycin treatment and then colonized by limited dilution. Single colonies were expanded for genomic DNA extraction and protein extraction. After expansion *Pcpe2^-/-^* cells were identified immunoblot analysis of for PCPE2 protein after SDS-PAGE of protein extracts.

### Culturing 3T3 Cells

Murine fibroblast 3T3-L1 cells were maintained in Dulbecco’s modified eagle medium (DMEM) supplemented with 10% calf bovine serum (CBS) following the manufacturer’s protocol.

### Differentiation of 3T3 Cells

Fully confluent low passage 3T3-L1 cells were kept in 10% CBS DMEM for 3 days, and then changed into 10% FBS DMEM supplemented with 1 µg/mL of insulin, 1 µM of dexamethasone, 0.5 mM of 3-isobutyl-1- methylxanthine and 1 µM of rosiglitazone. Two days later, cells were changed into 10% FBS DMEM with 1 µg/mL of insulin. After another 2 days cells were changed into 10% FBS DMEM and were continuously kept in this medium that was changed every 2 days. Differentiation of 3T3-L1 cells to adipocyte was monitored by visualizing the lipid droplets in the adipocytes by either oil red O staining of neutral lipids or by immunofluorescent staining of perilipin.

### Nondenaturing Gradient Gel Electrophoresis (33)

Aliquots were analyzed by 4–30% NDGGE to determine the particle diameter size homogeneity by comparison to high-molecular-weight standards (Amersham Biosciences) using either radioactivity or mass. Radioactive protein bands were visualized on the gel using a BAS 5000 phosphorimager (Fujifilm). Protein mass was quantified after staining with Simply Blue.

### Western Blotting

Western Blot was performed after total tissue/cell protein extraction using a radioimmune precipitation assay buffer (RIPA) (Cell Signaling Technologies, 9806S), or after detergent-free lipid raft isolation, or after co-immunoprecipitation (Co-IP). Aliquots of equal protein content (tissue homogenates or cell protein extracts) or equal volume (Co-IP eluents or lipid raft fractions) were diluted with 4X LDS buffer (Life Technologies, NP0007) to which solid DTT had been added to achieve a 100 mM final concentration. Samples were heated to 70 °C for 10 min, run on 12% SDS-PAGE, and then transferred to PVDF membrane (PerkinElmer, NEF1002001PK) using a semi-dry blot apparatus (Bio-Rad). Membranes were blocked with 5 % nonfat dry milk, incubated in primary antibodies (PCPE2 (Abcam, ab156224, 1:5,000), GAPDH (Ambion, AM4300, 1:20,000), Crk (BD, 610036, 1:2,000) and SR-BI (Novus, NB400-101, 1:5,000)) at 4 °C overnight, followed by horseradish peroxidase-conjugated secondary antibodies for 1 hour at room temperature. Finally blots were washed and incubated with chemiluminescence substrates, SuperSignal West Pico (Thermo Scientific, 34080) or Clarity (Bio-Rad, 170- 5060), and visualized with X-ray film. Band intensities were compared using Imge J software (NIH).

### Co-immunoprecipitation studies

Differentiated 3T3-L1 cells were washed twice with PBS (137 mM NaCl, 2.68 mM KCl, 10 mM Na_2_HPO_4_, pH 7.4) and lysed in a lysis buffer composed of 1 % NP-40 in 10 mM Tris pH 8.0, 150 mM NaCl, 2 mM EDTA, 2 mM EGTA, and several, different protease and phosphatase inhibitors. For adipose tissue 200 mg of tissue was homogenized in 500 µL of lysis buffer. Immunoprecipitation was accomplished by mixing 500 µg of protein extract with 500 µL of lysis buffer and then pre-cleared with 50 µL of 50% suspension of protein G agarose. After centrifugation the pre-cleared supernatant was incubated with anti-SR-BI antibody (Norvus Biologicals, NB400-101) overnight at 4°C. The next day 50 µL of a 50% suspension of protein G agarose was added and the mixture incubated at room temperature for 1 hour. The agarose beads were extensively washed and eluted first in soft elution buffer that contained 0.2 % (w/v) SDS, 0.1 % (v/v) Tween- 20, and 50 mM Tris-HCl (pH 8.0), which was followed by a hard elution using 2x NuPAGE LDS sample buffer (Life Technologies, NP0007).

### Rapid Cholesterol and Triglyceride Measurement

Plasma from both chow- and diet-fed mice was analyzed for total plasma and triglycerides by enzymatic assays, Wako Cholesterol E 999- 02601 and Thermo Fisher Scientific TR22421, respectively. Molecular species analysis of cholesteryl ester and triglyceride was performed using mass spectrometry.(33)

### Lipid Extraction

Lipids were extracted from tissue by heating in chloroform/methanol (2:1) overnight after adding 4 µg of [25,26,27-^13^C_3_]cholesterol as the internal standard for cholesterol analyses. A nonpolar lipid fraction containing cholesterol and cholesteryl esters was separated using the method of Kaluzny et al. (34) as previously reported (35). The eluate was evaporated under a stream of argon, dissolved in 1 ml of chloroform-methanol (1:1), and stored at −80°C until analyzed.

### Cholesterol quantitation by GC-MS

For FC analysis an aliquot of extract was evaporated under argon, dissolved in hexane, and then analyzed on a Thermo Scientific TSQ 8000 tandem mass spectrometer - Trace 1310 gas chromatograph (GC/MS/MS) equipped with a Triplus RSH autoinjector. The following parameters were employed: scan time 0.1 second, collision energy 10 V, emission current 25 µA, electron energy 42 eV, source and transfer line temperature 280° C, helium flow rate= 2.5E-8 m^3^/s, in the positive ion selective reaction monitoring mode: cholesterol, *m/z* 386.4→301.3; [25,26,27-^13^C_3_]cholesterol, *m/z* 389.9→304.3. Analysis was carried out using a TG-SQC column (15 m9 0.25 mm I.D.) with a 0.25-l m film thickness. For quantifying TC an equal aliquot was dried under a stream of nitrogen, dissolved in 1 mL of ethanol, mixed with 100 µL of 50% (w/w) aqueous potassium hydroxide, and then saponified for 1 hour at 65°C. After extraction TC was measured and CE calculated as the difference between FC and TC (35).

### Cholesteryl Ester Molecular Species

The tissue or plasma was extracted with cholesteryl heptadecanoate (Nu-Chek Prep) as the internal standard. An aliquot was dried down and dissolved in chloroform:methanol:24 mM aq ammonium formate (75:100:18.5, v/v/v). The solution was analyzed by direct infusion at 100 µl/min into an Agilent 6460 tandem mass spectrometer equipped with a JetStream™ interface. Analysis settings were: collision energy 15 V; capillary, 3000 V; nozzle 1000 V, source temp, 240°C; nitrogen was introduced at 7 l/min; nebulizing nitrogen was set to 18 l/min; collision gas, argon, at 1 mTorr. Precursors of *m/z* 369.1 were acquired with a dwell time of 200 ms.

### Triglyceride Analysis

Triheptadecanoylglycerol was added before extraction. An aliquot was dried down and dissolved in chloroform/300 mM ammonium formate in methanol (1:1.75, v/v). Each sample was infused at 100 µl/min, spray solvent chloroform/300 mM ammonium formate in methanol (1.23, v/v), into an Agilent 6460 tandem mass spectrometer equipped with a JetStream™ interface with positive ion mass spectra acquired from *m/z* 600 to 1,200. Triglycerides were quantified by comparison of the triglyceride peak intensities to triheptadecanoylglycerol. Analysis settings were: collision energy 15 V; capillary, 3000 V; nozzle 1000 V, source temp, 240°C; nitrogen was introduced at 7 l/min; nebulizing nitrogen was set to 18 l/min; dwell time 200 ms. To determine the distribution of fatty acids in TG neutral loss spectra were acquired for each of the major fatty acids.

## Acknowledgements

We would like to thank Dr. Ira J. Goldberg of the Department of Medicine, Columbia University College of Physicians and Surgeons for analyzing lipoprotein lipase activity.

## Footnotes

This work was supported by National Institute of Health Grants HL127649 (MST), HL58012 (DS), and American Heart Grant 14GRNT20500029 (MJT).

## Abbreviations

CBS: calf bovine serum
CHO: Chinese hampster overy
CE: cholesteryl ester
COE: cholesteryl-oleoyl-ether
Co-PI: co-immunoprecipitation
CUB: Complement C1r/C1s, Uegf, Bmp1
DMEM: Dulbecco’s modified eagle medium
FA: fatty acids
HDL: high density lipoprotein
FC: free cholesterol
LC-MS/MS: liquid chromatography-tandem mass spectrometry
LDLr: low density lipoprotein receptor
NDGGE: nondenaturing gradient gel electrophoresis
PCPE2: procollagen endopeptidase enhancer 2
PFO: perfluorooctanoic acid
SAT: subcutaneous fat
RIPA: radioimmune precipitation assay buffer
SR-BI: scavenger receptor class B1
TC: total cholesterol
VAT: visceral adipose tissue.

